# Required concentration index quantifies effective drug combinations against hepatitis C virus infection

**DOI:** 10.1101/2020.01.09.899807

**Authors:** Yusuke Kakizoe, Yoshiki Koizumi, Yukino Ikoma, Hirofumi Ohashi, Takaji Wakita, Shingo Iwami, Koichi Watashi

## Abstract

Successful clinical drug development requires rational design of combination treatments based on preclinical data. Anti-HCV drugs exhibit significant diversity in antiviral effect. Dose-response assessments can be used to determine parameters profiling the diverse antiviral effect during combination treatment. In the current study, a combined experimental and mathematical approaches were used to compare and score different combinations of anti-hepatitis C virus (HCV) treatments. A “required concentration index” was generated and used to rank the antiviral profile of possible double- and triple-drug combinations against HCV genotype 1 and 2. Rankings varied based on target HCV genotype. Interestingly, multidrug (double and triple) treatment not only augmented antiviral activity, but also reduced genotype-specific efficacy, suggesting another advantage of multidrug treatment. The current study provides a quantitative method for profiling drug combinations against viral genotypes, to better inform clinical drug development.

## INTRODUCTION

Newly approved antiviral drugs rely upon dosage, treatment period, and drug combinations established during clinical trials. Trials require large cohorts of patients, significant cost, extensive time and strict management of ethics and compliance: Different dose regimens, treatment times and drug combinations are evaluated during trials (1, 2). Additional trials are needed to establish drug efficacy against different viral genotypes (3-6). Despite the significant effort placed in clinical trials, escalation of dosage, increased treatment period, and combination therapy, significant improvement in efficacy have not always been realized.

Drug concentrations which achieve 50% virus reduction (*IC*_50_), can be used to characterize drug activity. Lower *IC*_50_ means that antiviral effects are achieved with lower concentrations of drug (7); however, a lower *IC*_50_ does not necessarily translate to higher antiviral effect. Antiviral effect depends on the Hill coefficient (*m*), in addition to *IC*_50_. A higher *m* value exponentially increases antiviral activity at higher doses (8-14). We have previously shown that *m* is unique to each anti-hepatitis C virus (HCV) drug, and that augmentation of antiviral activity with escalation of drug dose is quite diverse among the types of anti-HCV drugs (14). Multi-drug treatments also result in diverse effects depending on the drug combination. In-depth profiling of drug antiviral effects can be useful in designing a treatment protocol with maximal antiviral efficacy. Such profiling could result in significant savings in clinical trials. To date, antiviral efficacy variances between different anti-HCV drugs and drug combinations has not been characterized in detail.

HCV infection is a leading cause of liver cirrhosis and hepatocellular carcinoma, serious public health problems affecting approximately 170 million people worldwide (15). Recently, the development of new antiviral drugs known as direct acting antivirals (DAAs), have greatly improved treatment outcomes (16, 17). With rapid development, therapeutic DAA combinations have entered clinical practice without significant insight into the advantages and disadvantages of the various combinations. Further evaluation of HCV DAA effects could help identify the “best” available therapy and assist with optimizing combination treatments. A new quantitative method could also support evaluation of next generation anti-HCV treatments that could lead to the eradication of HCV. In the current study, we compare antiviral profiles of different classes of anti-HCV drugs to understand diversity of effects.

We recently developed a cell culture system combined with a mathematical model for quantifying anti-HCV drug efficacy at any concentration and multidrug combination (14). We systematically evaluated and compared the intrinsic anti-HCV activity of 15 antiviral agents and their combinations against HCV genotype 1. In the current study, we evaluate intrinsic anti-HCV activity in both genotype 1 and 2. We create an “effectiveness” ranking for HCV replication inhibition in mono- and multidrug cultures following exposure to high drug dose ranges. Significant diversity was observed between the antiviral activity profiles of different drugs. Thus, it is necessary to carefully select multidrug combinations to increase drug efficacy. We have demonstrated that the developed ranking index is able to delineate the advantages of past first-in-line anti-HCV treatment choices (14). Thus, in the current study, we use the combined cell culture plus mathematical modeling approach to quantify efficacy of diverse antiviral drug combinations. This framework could be applied to other diseases requiring multidrug treatment, such as tuberculosis and cancer.

## RESULTS

**Fig.1A** provides a schematic of the combined experimental and mathematical system that we previously developed for quantifying anti-HCV activity of drug(s) (14). In the previous study 14 anti-HCV agents were evaluated in mono and combination treatments against HCV genotype 1 (18). In the current study the same 14 drugs (**Table 1**) were tested against HCV genotype 1 (**Fig.1B**) and HCV genotype 2 (**Fig.1C**). Antiviral activity results from mono and combination treatments were used to develop a novel ranking index, the “required concentration index” or RCI (see below) (**MATERIALS** and **METHODS**). Note that 14 anti-HCV agents include 11 direct-acting antivirals (DAAs) including NS3 protease inhibitors [PIs; telaprevir (TPV), danoprevir (DPV), simeprevir (SMV), and asunaprevir (ASV)], a nucleoside NS5B polymerase inhibitor [NI; sofosbuvir (SOF)], non-nucleoside NS5B polymerase inhibitors [NNIs; VX-222 (VX), dasabuvir (DAS), nesbuvir (NSV), and tegobuvir (TGV)], and NS5A inhibitors [NS5AI; daclatasvir (DCV) and ledipasvir (LDV)] and 3 host-targeting agents (HTAs) included interferon-alpha (IFNα) and cyclophilin inhibitors [CIs; cyclosporin A (CsA) and SCY-635 (SCY)].

**Table 1.** Estimated characteristic parameters of the tested antiviral drugs

| Drug | Type | Target | Class | *IC*<sub>50</sub> |  | *m* |  | 435 |
| --- | --- | --- | --- | --- | --- | --- | --- | --- |
|  |  |  |  | genotype 1 | genotype 2 | genotype 1 | genotype 2 |  |
| TPV (nM) | DAA | NS3 protease | PI | 323.79 | – | 1.72 | – |  |
| DPV (nM) | DAA | NS3 protease | PI | 1.40 | – | 0.98 | – |  |
| SMV (nM) | DAA | NS3 protease | PI | 0.45 | 153.95 | 1.10 | 3.10 |  |
| ASV (nM) | DAA | NS3 protease | PI | 2.75 | 665.49 | 0.97 | 2.99 |  |
| SOF (nM) | DAA | NS5B polymerase NI | NI | 120.48 | 843.74 | 1.66 | 3.02 |  |
| VX (pM) | DAA | NS5B polymerase NNI | NNI | 107.58 | – | 1.81 | – |  |
| DAS (nM) | DAA | NS5B polymerase NNI | NNI | 1.50 | 7203.98 | 0.99 | 0.73 |  |
| NSV (nM) | DAA | NS5B polymerase NNI | NNI | 0.25 | – | 1.19 | – |  |
| TGV (nM) | DAA | NS5B polymerase NNI | NNI | 8.92 | – | 1.01 | – |  |
| DCV (nM) | DAA | NS5A | NS5AI | 0.10 | 0.13 | 1.11 | 3.68 |  |
| LDV (nM) | DAA | NS5A | NS5AI | 0.67 | 30.80 | 0.96 | 6.11 |  |
| IFNα (IU/ml) | HTA | – | IFN | 2.56 | – | 1.43 | – |  |
| CsA (μg/m) | HTA | Cyclophilin | CI | 0.40 | – | 1.53 | – |  |
| SCY (μM) | HTA | Cyclophilin | CI | 0.34 | – | 1.45 | – |  |

**Figure 1.**
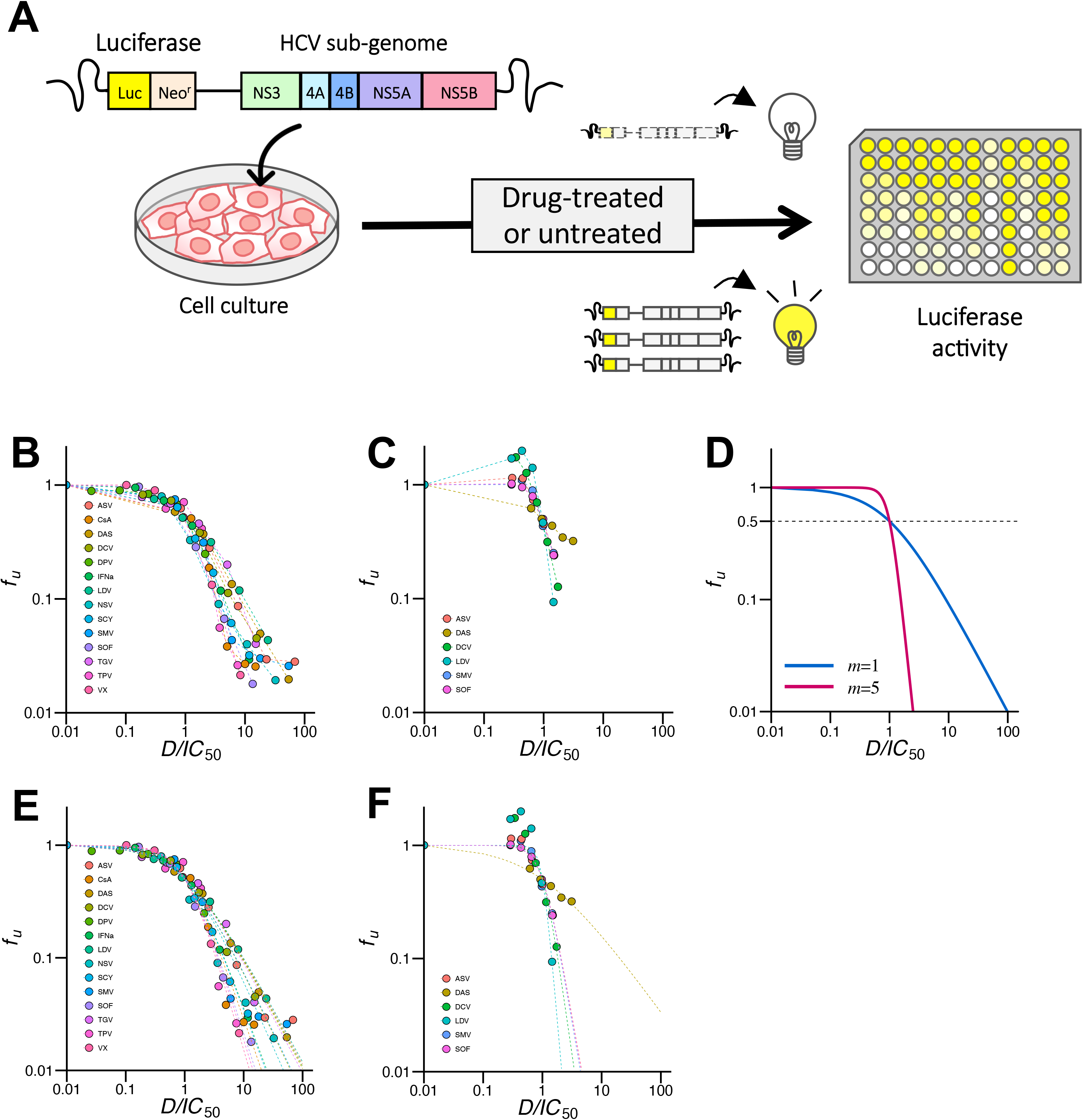
Experimental and mathematical system for quantifying antiviral activity of HCV drug(s): **(A)** Schematic representation of the method for quantifying HCV replication level. The assay uses HCV subgenomic replicons for genotype 1 and 2 carrying a fusion of the firefly luciferase gene (Luc) with or without the neomycin phosphotransferase (Neo^r^). The replicons autonomously and persistently replicate in Huh-7.5.1 cells (18, 28). Cells were incubated for 72 h with or without drug(s) then harvested and luciferase activity detected. Inhibition of HCV replication was quantified as the luciferase activity in drug-treated cells, relative to untreated cells. **(B)** Log–Log plots of dose-response curves normalized by *IC*_50_ (x-axis), determined from HCV genotype 1 subgenomic replicon assay of NS3 protease inhibitors (PIs; TPV, DPV, ASV, SMV), nucleoside-type NS5B polymerase inhibitor (NI; SOF), non-nucleoside-type NS5B polymerase inhibitors (NNIs; VX, DAS, NSV, TGV), NS5A inhibitors (NS5AIs; DCV, LDV), interferon (IFNα), and cyclophilin inhibitors (CIs; CsA, CSY). Each point represents the mean of three experiments. **(C)** Log–Log plots of dose-response curves from HCV genotype 2 subgenomic replicon assay of PIs (ASV, SMV), NI (SOF), NNIs (DAS), and NS5AIs (DCV, LDV). Each point represents the mean of three experiments. **(D)** Dose-response curves for hypothetical drugs with *m* = 1 and 5. Drugs with a higher *m* value show stronger antiviral activity at the same normalized drug concentration. **(E)** Least-square regression analysis was used to fit Eq.(1) to the corresponding dose-response curve for estimation of *IC*_50_ and *m* value for each drug against HCV genotype 1. **(F)** Least-square regression analysis was used to fit Eq.(1) to the corresponding dose-response curve for each drug against HCV genotype 2.

### Ranking anti-HCV mono-drug treatments

As shown in **Fig.1B** & **1C**, the antiviral profile of drugs against HCV genotypes 1 and 2 vary widely, suggesting that anti-HCV drugs exhibit strain-dependent effects. The typical dose-response curves of a single antiviral drug can be analyzed using the following hill function (14) (**Fig.1D**):

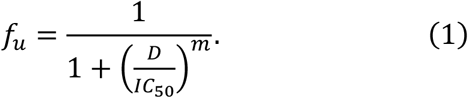

Here, *f_u_* represents the fraction of infection events unaffected by the drug (i.e., 1 − *f_u_* equals the fraction of drug-affected events). *D* is the drug concentration, *IC*_50_ is the drug concentration that achieves 50% inhibition of activity, and *m* is the slope of the dose-response curve (i.e., Hill coefficient) (14). Dose-response curves for drugs with higher *m* values show stronger antiviral activity at the same normalized drug concentration so long as the drug concentration is higher than *IC*_50_ (**Fig.1D**). Least-square regression analysis was used to fit Eq.(1) to dose-response curves (**Fig.1E** & **1F**) and estimate *IC*_50_ and *m* values. Estimated values for each drug against each HCV genotype are summarized in **Table 1**. The hill function may not accurately fit the dose-response curve at lower drug concentrations (**Fig.1F**, especially for doses lower than *IC*_50_). Typical clinical drug concentrations are around 10- to 100-fold of *IC_50_,* therefore it is generally possible to quantify effectiveness of anti-HCV drug(s) with this method especially for such a high drug concentrations. As discussed in recent publications (8-14), both *IC*_50_ and *m* values are needed to accurately estimate antiviral drug potency, though only *IC*_50_ is widely used in the drug development field. Since estimated values for each drug differ relative to target HCV genotype, it is important to optimize mono and combination therapy against each genotype.

To characterize efficacy of drugs, we calculated a “required concentration index” (RCI) for each anti-HCV drug against genotype 1 and 2. The RCI represents the critical fold increase of *IC*_50_ requiring 1 − *f_u_* inhibition of viral replication.

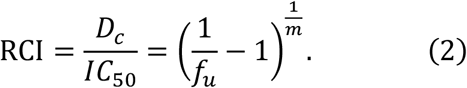

Here, *D_c_* is the critical drug concentration satisfying Eq.(2). Drugs with small RCI values are more efficient inhibitors of HCV replication than drugs with high RCI. Interestingly, high *m* tends to be associated with smaller RCI. By substituting estimated *IC*_50_ and *m* parameters and setting *f_u_* to 0.05 in Eq.(2), we calculated the RCI required for 95% inhibition of HCV replication (i.e., RCI_95_). We summarize RCI_95_ values of each drug against genotypes 1 and 2 in **Fig.2A** & **2B**, respectively. It should be noted that SOF, a nucleoside-type polymerase inhibitor used as a key agent in current and past DAA combinations, was effective in both genotype 1 and 2, which is consistent with SOF’s known clinical pan-genotypic anti-HCV characteristic (19).

**Figure 2.**
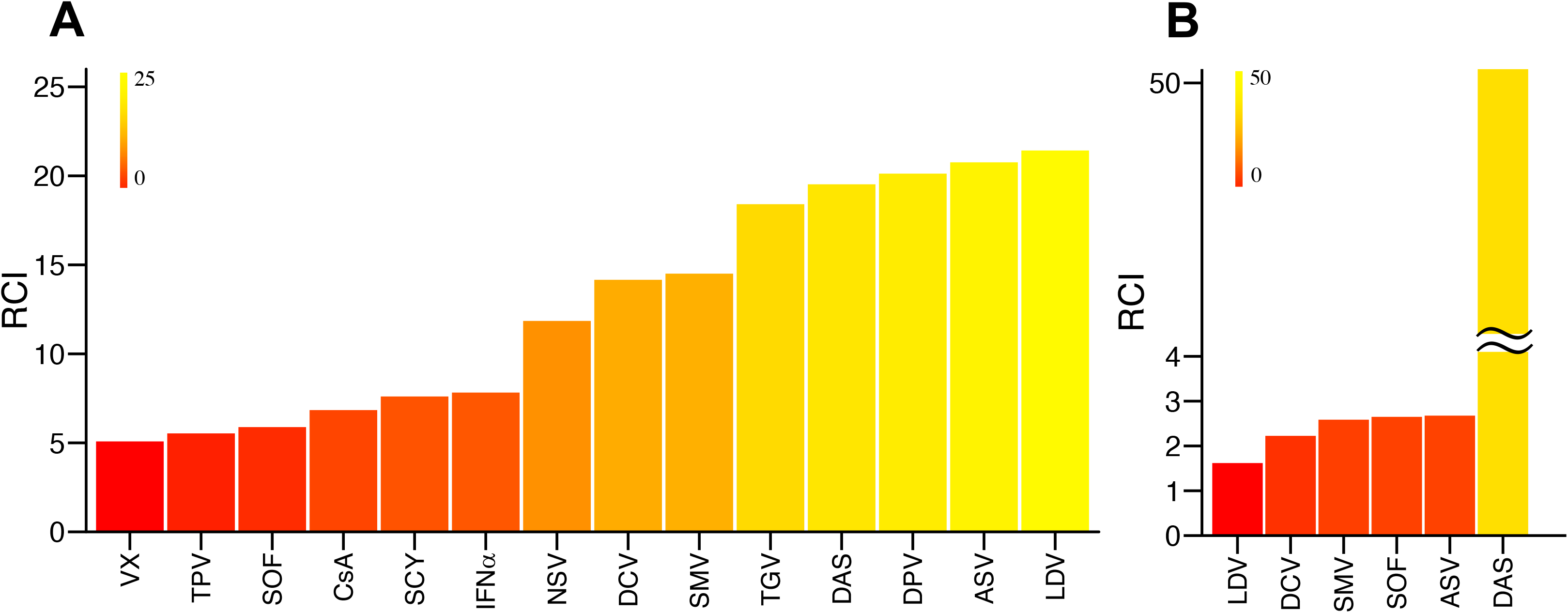
Ranking anti-HCV mono drug treatments against genotypes 1 and 2: The critical dose of antiviral drug (i.e., fold increase of *IC*_50_) required to inhibit viral replication by 95%, i.e., RCI95, was calculated for HCV **(A)** genotype 1 and **(B)** genotype 2.

### Ranking anti-HCV multi-drug treatments

Using the replicon system, the antiviral activity of double- and triple-drug combinations (**Fig.3** & **Fig.4**) were investigated using consistent ratios of drug concentrations (i.e., 0.25 × *IC*_50_, 0.5 × *IC_50_, 1 × *IC*_50_, 2 × *IC*_50_,* and 4 × *IC*_50_).

**Figure 3.**
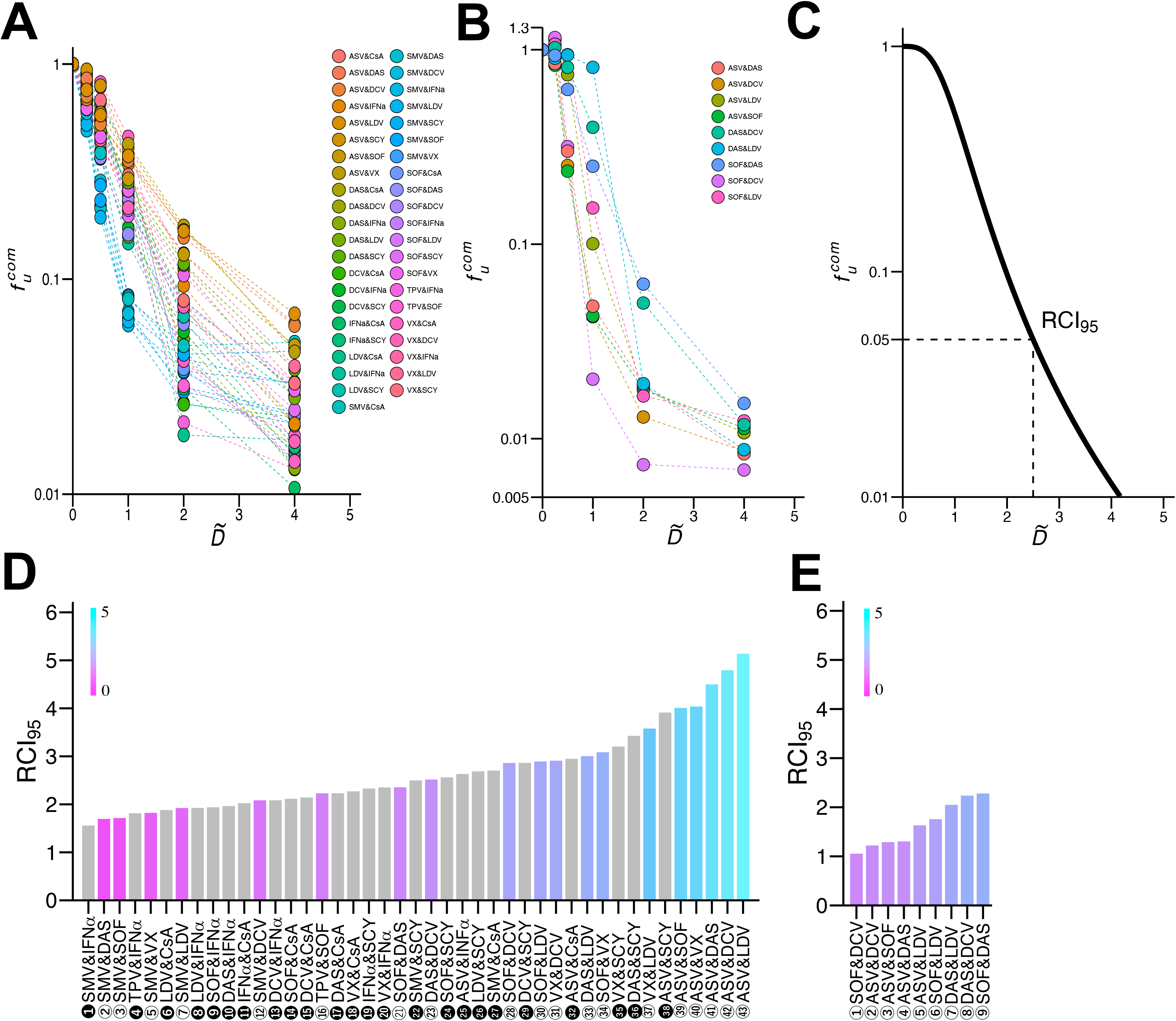
Quantification of anti-HCV double-drug combinations against genotype 1 and 2: Log–log plots of dose-response curves for 43 **(A)** and 9 **(B)** double-drug combinations of inter-class (or subclass) antiviral drugs against genotype 1 **(A)** and genotype 2 **(B)**. Each point represents the mean of four experiments. For comparison of drugs, concentration of each drug was based on a consistent ratio of its *IC*_50_ value. **(C)** Dose-response curve for a hypothetical double-drug combination. RCI_95_ for (double or triple-) drug combinations can be determined from the point where the curve intersects 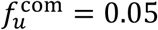 (dashed line). **(D)** and **(E)** show RCI_95_ rankings for 43 double-combinations against HCV genotype 1 and 9 double-combinations against HCV genotype 2, respectively. Combinations with gray bars and black numbers (e.g., 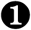) correspond to treatment regimens that include HTAs. Light pink to blue bars and white numbers (e.g., 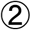) correspond to treatment combinations with DAA-only.

**Figure 4.**
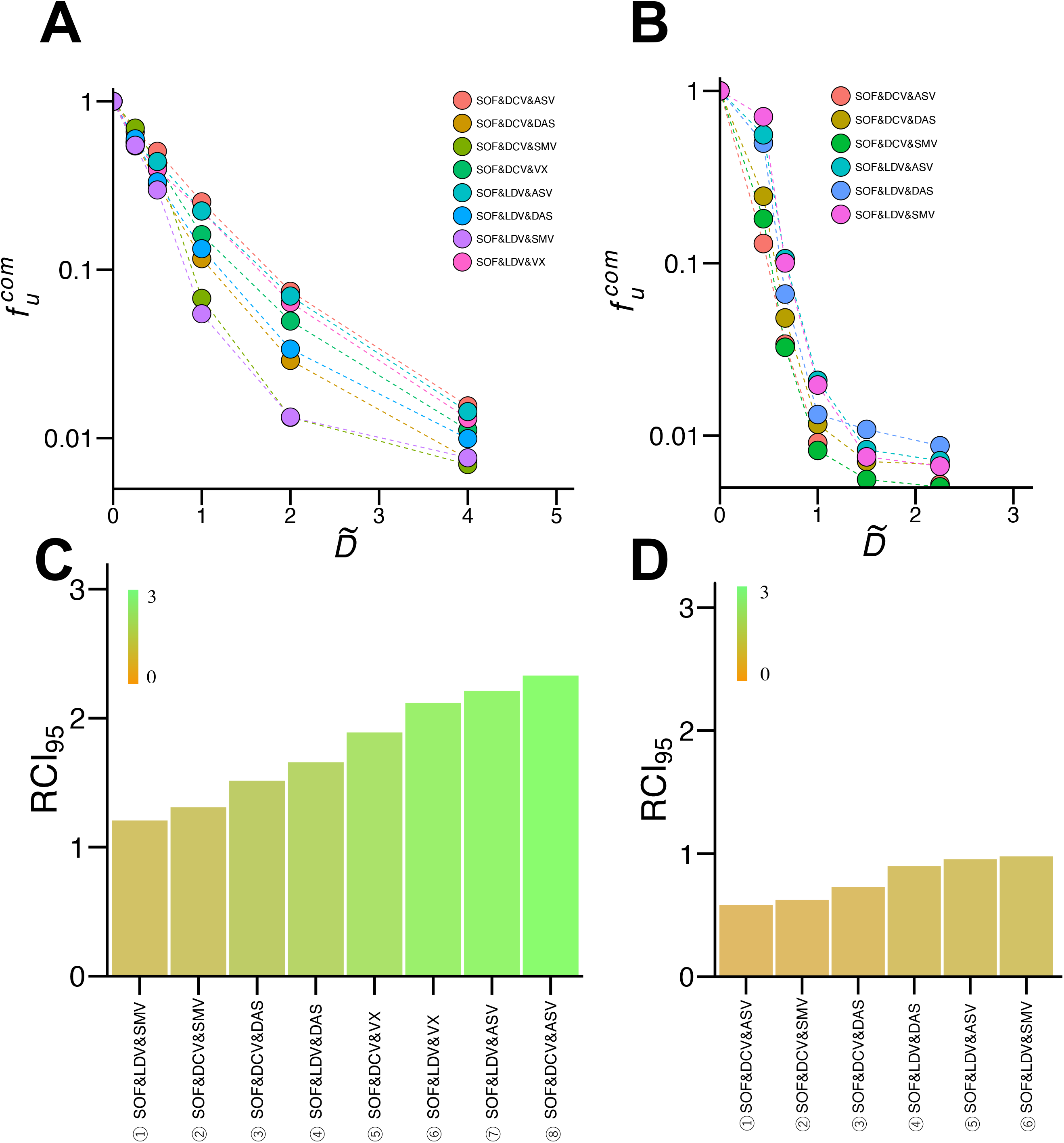
Quantification of anti-HCV triple-drug combinations against genotype 1 and 2: Log–log plots of dose-response curves for 8 **(A)** and 6 **(B)** triple-drug combinations against HCV genotype 1 **(A)** and 2 **(B)**. Each point represents the mean of five experiments for genotype 1 and four experiments for genotype 2. For comparison of drugs, concentration of each drug was based on a consistent ratio of its *IC*_50_ value. **(C)** and **(D)** show RCI_95_ rankings of the triple-drug combinations against HCV genotype 1 and 2, respectively.

Inhibitory activity was evaluated for 43 double drug combinations against HCV genotype 1, and, 9 double drug combinations against genotype 2. Results are shown in **Fig.3A** & **3B**, respectively. Here, *D_a_, D_b_, …, D_i_* are defined as the concentration of drug a, b, …, i and 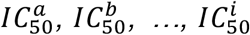 refer to the corresponding *IC*_50_. Combined drug concentration in these experiments is described as 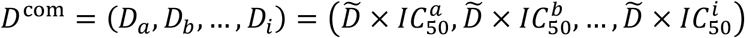, where 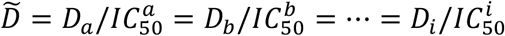 is the constant ratio to *IC*_50_ of each combined drug (x-axis of dose-response curves). As shown in **Fig.3C**, a similar hill function can be fit to dose-response curves of drug combinations (14):

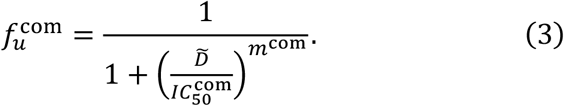

Here, 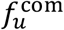 is the fraction of infection events unaffected by the drug combination, 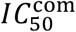 is the constant ratio that inhibits HCV replication by 50%, and *m*^com^ is the Hill coefficient (14). In **Table 2**, we summarize estimated parameters, 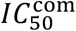 and *m*^com^, for double-drug combinations.

**Table 2.**
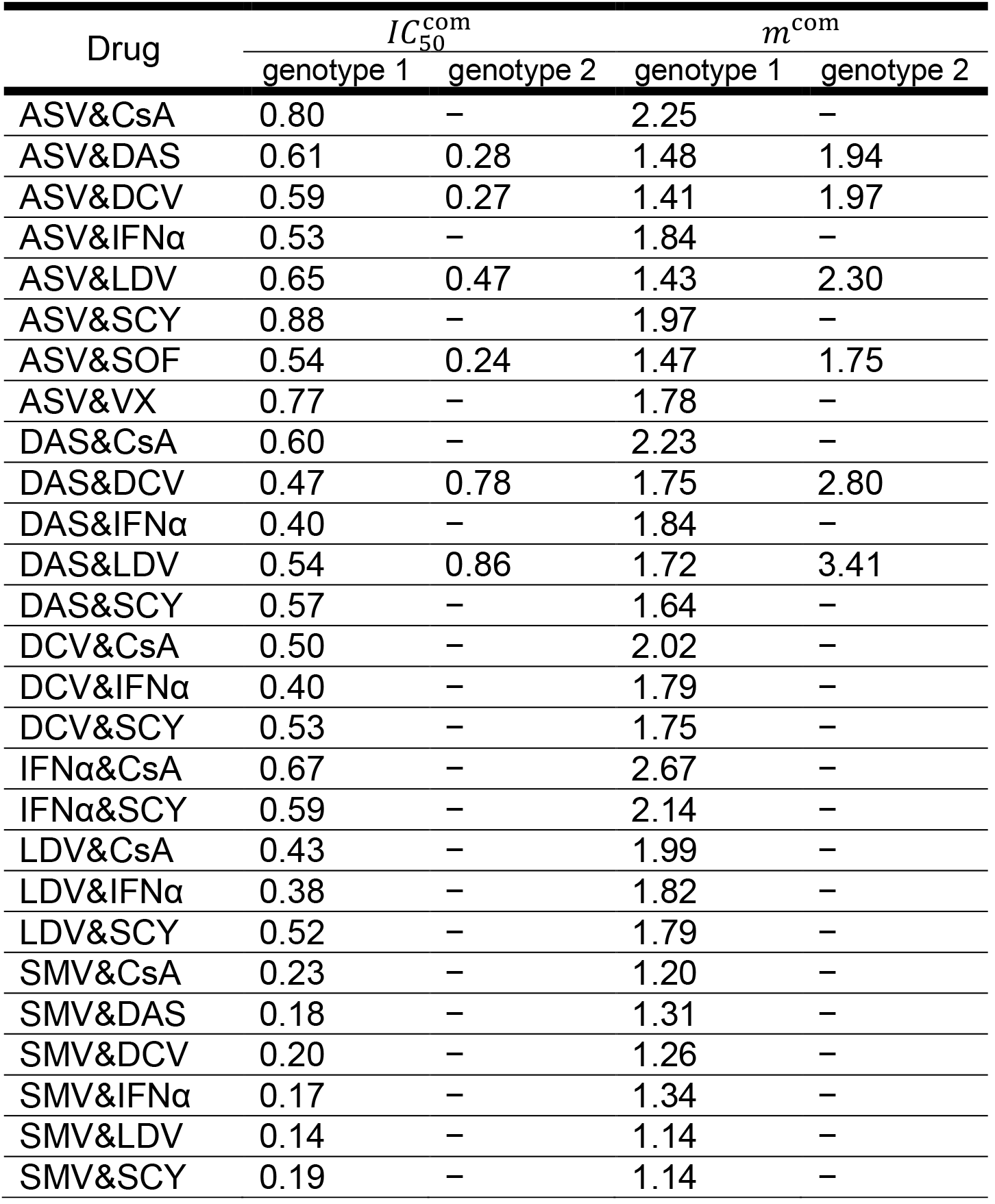
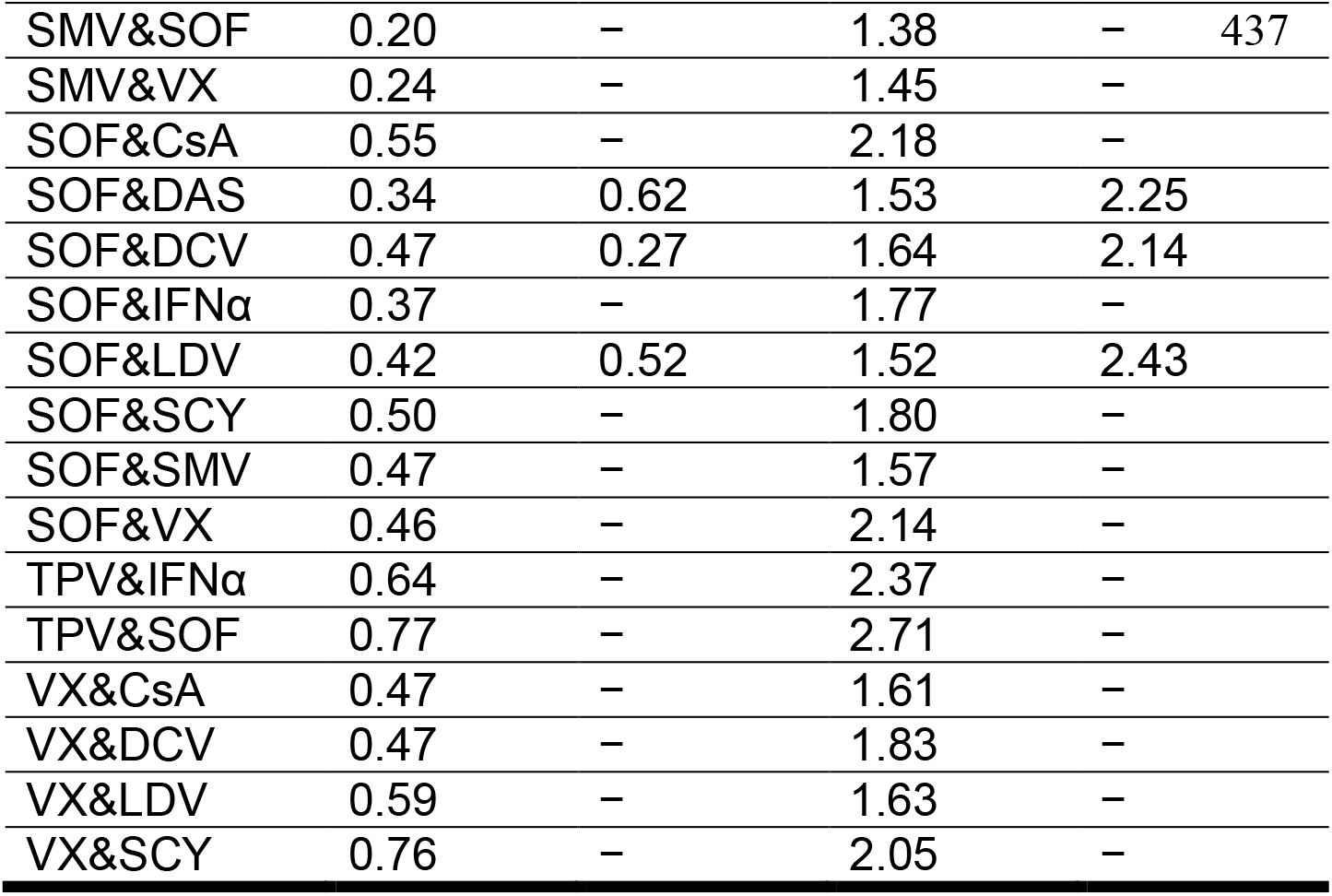
Estimated characteristic parameters of the antiviral drug combinations

| Drug | $IC_{50}^{com}$ | | $m^{com}$ | |
| --- | --- | --- | --- | --- |
|  | genotype 1 | genotype 2 | genotype 1 | genotype 2 |
| ASV&CsA | 0.80 | – | 2.25 | – |
| ASV&DAS | 0.61 | 0.28 | 1.48 | 1.94 |
| ASV&DCV | 0.59 | 0.27 | 1.41 | 1.97 |
| ASV&IFNα | 0.53 | – | 1.84 | – |
| ASV&LDV | 0.65 | 0.47 | 1.43 | 2.30 |
| ASV&SCY | 0.88 | – | 1.97 | – |
| ASV&SOF | 0.54 | 0.24 | 1.47 | 1.75 |
| ASV&VX | 0.77 | – | 1.78 | – |
| DAS&CsA | 0.60 | – | 2.23 | – |
| DAS&DCV | 0.47 | 0.78 | 1.75 | 2.80 |
| DAS&IFNα | 0.40 | – | 1.84 | – |
| DAS&LDV | 0.54 | 0.86 | 1.72 | 3.41 |
| DAS&SCY | 0.57 | – | 1.64 | – |
| DCV&CsA | 0.50 | – | 2.02 | – |
| DCV&IFNα | 0.40 | – | 1.79 | – |
| DCV&SCY | 0.53 | – | 1.75 | – |
| IFNα&CsA | 0.67 | – | 2.67 | – |
| IFNα&SCY | 0.59 | – | 2.14 | – |
| LDV&CsA | 0.43 | – | 1.99 | – |
| LDV&IFNα | 0.38 | – | 1.82 | – |
| LDV&SCY | 0.52 | – | 1.79 | – |
| SMV&CsA | 0.23 | – | 1.20 | – |
| SMV&DAS | 0.18 | – | 1.31 | – |
| SMV&DCV | 0.20 | – | 1.26 | – |
| SMV&IFNα | 0.17 | – | 1.34 | – |
| SMV&LDV | 0.14 | – | 1.14 | – |
| SMV&SCY | 0.19 | – | 1.14 | – |

|  |  |  |  |  |  |
| --- | --- | --- | --- | --- | --- |
| SMV&SOF | 0.20 | – | 1.38 | – | 437 |
| SMV&VX | 0.24 | – | 1.45 | – |  |
| SOF&CsA | 0.55 | – | 2.18 | – |  |
| SOF&DAS | 0.34 | 0.62 | 1.53 | 2.25 |  |
| SOF&DCV | 0.47 | 0.27 | 1.64 | 2.14 |  |
| SOF&IFNα | 0.37 | – | 1.77 | – | |
| SOF&LDV | 0.42 | 0.52 | 1.52 | 2.43 |  |
| SOF&SCY | 0.50 | – | 1.80 | – |  |
| SOF&SMV | 0.47 | – | 1.57 | – |  |
| SOF&VX | 0.46 | – | 2.14 | – |  |
| TPV&IFNα | 0.64 | – | 2.37 | – | |
| TPV&SOF | 0.77 | – | 2.71 | – |  |
| VX&CsA | 0.47 | – | 1.61 | – |  |
| VX&DCV | 0.47 | – | 1.83 | – |  |
| VX&LDV | 0.59 | – | 1.63 | – |  |
| VX&SCY | 0.76 | – | 2.05 | – |  |

Similar to mono treatments, the required concentration index for drug combinations is derived as

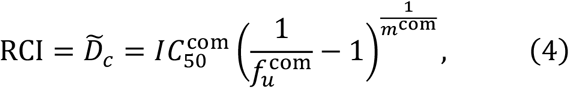

The RCI required for 95% inhibition of HCV replication is extrapolated from the point at which the curve intersects 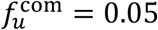 (dashed line in **Fig.3C**). Note that the critical constant ratio, 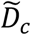, satisfying Eq.(4) can be uniquely determined. The RCI_95_ values for double-drug combinations against genotype 1 and 2 are summarized in **Fig.3D** & **3E**, respectively. RCI_95_ varies depending on drug combination. For genotype 1, RCI_95_ ranged from 1.56 to 5.14, for genotype 2 RCI_95_ ranged from 1.05 to 2.28. The drug combination with the best anti-HCV profile against genotype 1 is SMV plus IFNα; **Fig.3D**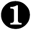. This combination used to be the first-in-line anti-HCV drug prior to the development of DAA treatments (17). Combinations including a non-DAA are presented as gray bars with black number designations. Combinations with DAA-only double treatments are plotted in light pink to blue and designated with white numbers (**Fig.3D**). For the DAA-only combinations, one of the most effective treatments against genotype 1 was the combination of SMV and SOF (**Fig.3D**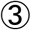), a primary treatment choice in the early era of DAA-only treatment (16). A long term first-in-line DAA combination, SOF and LDV (**Fig.3D**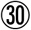 & **3E**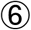), ranked in the midrange of efficacy against both genotype 1 and 2. Most other drug combinations ranked differently against genotype 1 and genotype 2. ASV plus LDV (**Fig.3D**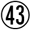 & **3E**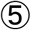) was the least effective DAA-only combination against genotype 1, but fell in the mid-range for effectiveness against genotype 2. SOF plus DAS (**Fig.3D**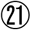 & **3E**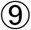) ranked in the mid-range against genotype 1, but ranked lowest against genotype 2. These trends suggest an overall difference in drug effect depending on the target HCV genotype, and indicate the importance of profiling drugs against each genotype.

Eight triple-DAA treatments were profiled against HCV genotype 1 and 6 triple-combinations were evaluated against genotype 2 (**Fig.4A** & **4B**). Triple combination assessments included NS3 protease inhibitor (SMV, ASV) with NS5A inhibitor (DCV, LDV) and NI NS5B polymerase inhibitor (SOF), or NS5A inhibitor with NI NS5B polymerase inhibitor and NNI NS5B polymerase inhibitor (VX, DAS). 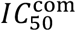 for triple-drug combinations are summarized in **Table 3**. RCI_95_ values of triple-drug combinations against genotype 1 and 2 are summarized in **Fig.4C** & **4D**, respectively. RCI_95_ values ranged from 1.21 to 2.33 for genotype 1 and 0.58 to 0.98 for genotype 2. Triple combination treatment with SOF, LDV and SMV was most effective against genotype 1 (**Fig.4C**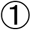), and least effective against genotype 2 (**Fig.4D**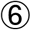). SOF plus DCV and SMV (**Fig.4C**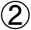) was also significantly effective against genotype 1, consistent with the reported clinical efficacy of this triple combination (20, 21).

**Table 3.** Estimated characteristic parameters of the antiviral drug combinations

| Drug | $IC_{50}^{com}$ | | $m^{com}$ | |
| --- | --- | --- | --- | --- |
|  | genotype 1 | genotype 2 | genotype 1 | genotype 2 |
| SOF&DCV&VX | 0.40 | – | 1.92 | – |
| SOF&DCV&ASV | 0.55 | 0.13 | 2.03 | 2.09 |
| SOF&DCV&DAS | 0.34 | 0.20 | 1.97 | 2.28 |
| SOF&DCV&SMV | 0.33 | 0.16 | 2.14 | 2.23 |
| SOF&LDV&VX | 0.45 | – | 1.90 | – |
| SOF&LDV&ASV | 0.46 | 0.33 | 1.88 | 2.88 |
| SOF&LDV&DAS | 0.35 | 0.26 | 1.88 | 2.49 |
| SOF&LDV&SMV | 0.26 | 0.35 | 1.90 | 3.04 |

The correlation in ranking between the required concentration index and clinical data suggest that this method could assist with the search for drugs that achieve an efficient antiviral inhibition with different HCV genotypes.

## DISCUSSION

Our study shows that the concentration of drug (calculated as fold of *IC*_50_), that achieves 95% virus inhibition (RCI95), highly varied depending on the type of drug and combination with other drugs. RCI95 of drugs in mono treatment ranged as much as 4.2 fold in antiviral activity against HCV genotype 1 (**Fig.2A**, RCI_95_=5.08-21.4). This diversity in RCI95 indicates the importance of characterizing more than just the IC_50_ of drugs when predicting antiviral efficacy in clinical settings. In double-drug combinations, RCI_95_ values decreased (**Fig.3D** & **3E**) compared with mono treatments (**Fig.2A** & **2B**), indicating elevated antiviral activity resulted from combination treatment. The RCI_95_ values of DAA-only double combinations ranged from 1.70 (SMV & DAS) to 5.14 (ASV & LDV) in genotype 1 and from 1.05 (SOF & DCV) to 2.28 (SOF & DAS) in genotype 2. Thus, the diversity in RCI95 is different among genotypes. Genotype differences are probably due differences in replication activity and the varied dependency on target (22, 23).

Triple DAA treatments have become the final strategy for improving treatment outcomes, especially with difficult-to-treat HCV. Triple combinations are also used as a means to shorten treatment periods. Understanding the activity of triple DAA combinations is important in advancing towards worldwide eradication of HCV virus (24-27). Consistent with ongoing clinical trials which show higher treatment efficacy of triple-drug combinations, triple combinations reduced RCI95 beyond double-drug combination levels (**Fig.4C** & **4D**). RCI_95_ for triple drug combinations ranged from 1.21 to 2.33 in genotype 1 and from 0.58 to 0.98 in genotype 2. Interestingly, the RCI_95_ values of selected drugs (SMV, ASV, DCV, LDV, DAS, VX and SOF) were less variable in triple-drug combinations compared with double combinations. RCI95 ranged 1.9 fold (1.21 for SOF & LDV & SMV to 2.33 for SOF & DCV & ASV) with triple-drug combinations, 3.0 fold (1.70 for SMV & DAS to 5.14 for ASV & LDV) in double-drug combinations, and 4.2 fold (5.08 for VX to 21.4 for LDV) in single-drug treatment against genotype 1. These data suggest that multidrug treatments such as triple-drug combinations provide more consistent antiviral effect irrespective of the choice of drugs, yet another advantage of triple combinations.

In an era of rapidly progressing anti-HCV treatments, selection of the “best” combination treatment is critical to establishing the next generation of anti-HCV treatments against difficult-to-treat HCV and eventually eradicating HCV. We have developed an integrated experimental and mathematical method to evaluate the efficacy of anti-HCV dugs against HCV genotype 1 and 2. The method was used to score mono- and multi-drug treatment regimens against HCV. This scoring could be used to optimize multidrug treatment regimens prior to clinical entry.

## METHODS

Anti-HCV effect of each drug against genotype 1b or genotype 2a was evaluated with subgenomic replicon systems. As a genotype 1b model, LucNeo#2 (LN2) cells were employed that carry a dicistronic subgenomic replicon including open reading frames (ORFs) for the firefly luciferase-neomycin phosphotransferase fusion protein (translated by HCV 5’-untranslated region) and the NS3–NS5B region of HCV genotype 1b strain NN (translated by encephalomyocarditis virus (EMCV) internal ribosome entry site) (18). Huh-7.5.1 cells transfected with a subgenomic replicon that included the ORFs for the NS3–NS5B region of HCV genotype 2a strain JFH-1 and the firefly luciferase gene (SGR-JFH1/Luc) were used for a genotype 2a model (28). These cells were seeded at 7 × 10^3^ cells per well and treated with indicated concentrations of various drugs. Following 72 h of incubation, cells were lysed and cellular luciferase activity was measured to evaluate the HCV replication activity with a Luciferase Assay System (Promega) per manufacturer’s protocol (18).

Fourteen anti-HCV drugs were evaluated as single treatments. Eleven of these were direct-acting antivirals (DAAs) of the following classes: NS3/4A protease inhibitors [PIs: telaprevir (TPV), danoprevir (DPV), simeprevir (SMV), and asunaprevir (ASV)], nucleoside NS5B polymerase inhibitor [NI: sofosbuvir (SOF)], non-nucleoside NS5B polymerase inhibitors [NNIs: VX-222 (VX), dasabuvir (DAS), nesbuvir (NSV), and tegobuvir (TGV)], and NS5A inhibitors [NS5AI: daclatasvir (DCV) and ledipasvir (LDV)]. The other 3 drugs tested were host-targeting agents (HTAs) including interferon-alpha (IFNα) and cyclophilin inhibitors [Cis: cyclosporin A (CsA) and SCY-635. For multidrug studies, cells were treated with combinations of two or three drugs prior to evaluation of activity. All anti-HCV agents were purchased or kindly provided as described (14).

## ACKNOWLEDGMENTS

We are grateful to Dr. Kunitada Shimotohno at National Center for Global Health and Medicine for providing LucNeo#2 cells. SCY635 was kindly provided by Scynexis, Inc. We appreciate the editorial assistance provided by Dr. Senko Tsukuda at Department of Virology II, National Institute of Infectious. This study was supported in part by Grants-in-Aid for JSPS Scientific Research (KAKENHI) Scientific Research B 18KT0018 (to S.I.), 18H01139 (to S.I.), 16H04845 (to S.I.), 17H04085 (to K.W.), Scientific Research in Innovative Areas 19H04839 (to S.I.), 18H05103 (to S.I.); AMED CREST 19gm1310002 (to S.I.); AMED J-PRIDE 19fm0208006s0103 (to S.I.), 19fm0208014h0003 (to S.I.), 19fm0208019h0103 (to S.I.), 19fm0208019j0003 (to K.W.); AMED Research Program on HIV/AIDS 19fk0410023s0101 (to S.I.); Research Program on Emerging and Re-emerging Infectious Diseases 19fk0108050h0003 (to S.I.); Program for Basic and Clinical Research on Hepatitis 19fk0210036h0502 (to S.I.), 19fk0210036j0002 (to K.W.); Program on the Innovative Development and the Application of New Drugs for Hepatitis B 19fk0310114h0103 (to S.I.), 19fk0310114j0003 (to K.W.), 19fk0310101j1003 (to K.W.), 19fk0310103j0203 (to K.W.); JST MIRAI (to S.I. and K.W.); JST CREST (to S.I. and K.W.); Mitsui Life Social Welfare Foundation (to S.I. and K.W.); Shin-Nihon of Advanced Medical Research (to S.I.); Suzuken Memorial Foundation (to S.I.); Life Science Foundation of Japan (to S.I.); SECOM Science and Technology Foundation (to S.I.); The Japan Prize Foundation (to S.I.); Toyota Physical and Chemical Research Institute (to S.I.); Fukuoka Financial Group, Inc. (to S.I.); Kyusyu Industrial Advancement Center Gapfund Program (to S.I.); Foundation for the Fusion Of Science and Technology (to S.I.); The Yasuda Medical Foundation (to K.W.); Smoking Research Foundation (to K.W.); Takeda Science Foundation (to K.W.); Mochida Memorial Foundation for Medical and Pharmaceutical Research (to K.W.).

## AUTHOR CONTRIBUTIONS

SI and KW designed the experiments. HO and KW conducted the experiments. YK, YK and YI carried out the computational analyses. SI and KW supervised the project. All authors contributed to the manuscript text.

## References

1. Pawlotsky JM. New hepatitis C therapies: the toolbox, strategies, and challenges. Gastroenterology. 2014;146(5):1176–92.

2. Schinazi R, Halfon P, Marcellin P, and Asselah T. HCV direct-acting antiviral agents: the best interferon-free combinations. Liver Int. 2014;34 Suppl 1:69–78.

3. Afdhal N, Zeuzem S, Kwo P, Chojkier M, Gitlin N, Puoti M, et al. Ledipasvir and sofosbuvir for untreated HCV genotype 1 infection. N Engl J Med. 2014;370(20):1889–98.

4. Gane EJ, Hyland RH, Yang Y, Svarovskaia E, Stamm LM, Brainard DM, et al. Efficacy of Ledipasvir Plus Sofosbuvir for 8 or 12 Weeks in Patients With Hepatitis C Virus Genotype 2 Infection. Gastroenterology. 2017;152(6):1366–71.

5. Gane EJ, Hyland RH, An D, Svarovskaia E, Pang PS, Brainard D, et al. Efficacy of ledipasvir and sofosbuvir, with or without ribavirin, for 12 weeks in patients with HCV genotype 3 or 6 infection. Gastroenterology. 2015;149(6):1454–61.e1.

6. Kohli A, Kapoor R, Sims Z, Nelson A, Sidharthan S, Lam B, et al. Ledipasvir and sofosbuvir for hepatitis C genotype 4: a proof-of-concept, single-centre, open-label phase 2a cohort study. Lancet Infect Dis. 2015;15(9):1049–54.

7. Aykul S, and Martinez-Hackert E. Determination of half-maximal inhibitory concentration using biosensor-based protein interaction analysis. Anal Biochem. 2016;508:97–103.

8. Shen L, Peterson S, Sedaghat AR, McMahon MA, Callender M, Zhang H, et al. Dose-response curve slope sets class-specific limits on inhibitory potential of anti-HIV drugs. Nat Med. 2008;14(7):762–6.

9. Sampah ME, Shen L, Jilek BL, and Siliciano RF. Dose-response curve slope is a missing dimension in the analysis of HIV-1 drug resistance. Proc Natl Acad Sci U S A. 2011;108(18):7613–8.

10. Shen L, Rabi SA, and Siliciano RF. A novel method for determining the inhibitory potential of anti-HIV drugs. Trends Pharmacol Sci. 2009;30(12):610–6.

11. Jilek BL, Zarr M, Sampah ME, Rabi SA, Bullen CK, Lai J, et al. A quantitative basis for antiretroviral therapy for HIV-1 infection. Nat Med. 2012;18(3):446–51.

12. Shen L, Rabi SA, Sedaghat AR, Shan L, Lai J, Xing S, et al. A critical subset model provides a conceptual basis for the high antiviral activity of major HIV drugs. Sci TranslMed. 2011;3(91):91ra63.

13. Laskey SB, and Siliciano RF. A mechanistic theory to explain the efficacy of antiretroviral therapy. Nat Rev Microbiol. 2014;12(11):772–80.

14. Koizumi Y, Ohashi H, Nakajima S, Tanaka Y, Wakita T, Perelson AS, et al. Quantifying antiviral activity optimizes drug combinations against hepatitis C virus infection. Proc Natl Acad Sci U S A. 2017;114(8):1922–7.

15. Lauer GM, and Walker BD. Hepatitis C virus infection. N Engl J Med. 2001;345(1):41–52.

16. Pawlotsky JM. Hepatitis C treatment: the data flood goes on-an update from the liver meeting 2014. Gastroenterology. 2015;148(3):468–79.

17. Pawlotsky JM, Feld JJ, Zeuzem S, and Hoofnagle JH. From non-A, non-B hepatitis to hepatitis C virus cure. J Hepatol. 2015;62(1 Suppl):S87–99.

18. Goto K, Watashi K, Murata T, Hishiki T, Hijikata M, and Shimotohno K. Evaluation of the anti-hepatitis C virus effects of cyclophilin inhibitors, cyclosporin A, and NIM811. Biochem Biophys Res Commun. 2006;343(3):879–84.

19. Feld JJ, Jacobson IM, Hezode C, Asselah T, Ruane PJ, Gruener N, et al. Sofosbuvir and Velpatasvir for HCV Genotype 1, 2, 4, 5, and 6 Infection. N Engl J Med. 2015;373(27):2599–607.

20. Lau G, Benhamou Y, Chen G, Li J, Shao Q, Ji D, et al. Efficacy and safety of 3-week response-guided triple direct-acting antiviral therapy for chronic hepatitis C infection:a phase 2, open-label, proof-of-concept study. Lancet Gastroenterol Hepatol. 2016;1(2):97–104.

21. Sulkowski MS, Feld JJ, Lawitz E, Felizarta F, Corregidor AM, Khalid O, et al. Efficacy and safety of 6 or 8 weeks of simeprevir, daclatasvir, sofosbuvir for HCV genotype 1 infection. J Viral Hepat. 2018;25(6):631–9.

22. Scheel TK, Gottwein JM, Mikkelsen LS, Jensen TB, and Bukh J. Recombinant HCV variants with NS5A from genotypes 1-7 have different sensitivities to an NS5A inhibitor but not interferon-alpha. Gastroenterology. 2011;140(3):1032–42.

23. McPhee F, Sheaffer AK, Friborg J, Hernandez D, Falk P, Zhai G, et al. Preclinical Profile and Characterization of the Hepatitis C Virus NS3 Protease Inhibitor Asunaprevir (BMS-650032). Antimicrob Agents Chemother. 2012;56(10):5387–96.

24. Kohli A, Osinusi A, Sims Z, Nelson A, Meissner EG, Barrett LL, et al. Virological response after 6 week triple-drug regimens for hepatitis C: a proof-of-concept phase 2A cohort study. Lancet. 2015;385(9973):1107–13.

25. Everson GT, Sims KD, Rodriguez-Torres M, Hezode C, Lawitz E, Bourliere M, et al. Efficacy of an interferon- and ribavirin-free regimen of daclatasvir, asunaprevir, and BMS-791325 in treatment-naive patients with HCV genotype 1 infection. Gastroenterology. 2014;146(2):420–9.

26. Poordad F, Sievert W, Mollison L, Bennett M, Tse E, Brau N, et al. Fixed-dose combination therapy with daclatasvir, asunaprevir, and beclabuvir for noncirrhotic patients with HCV genotype 1 infection. Jama. 2015;313(17):1728–35.

27. Muir AJ, Poordad F, Lalezari J, Everson G, Dore GJ, Herring R, et al. Daclatasvir in combination with asunaprevir and beclabuvir for hepatitis C virus genotype 1 infection with compensated cirrhosis. Jama. 2015;313(17):1736–44.

28. Kato T, Date T, Miyamoto M, Sugiyama M, Tanaka Y, Orito E, et al. Detection of anti-hepatitis C virus effects of interferon and ribavirin by a sensitive replicon system. J Clin Microbiol. 2005;43(11):5679–84.

